# Crossbill: an open access single objective light-sheet microscopy platform

**DOI:** 10.1101/2021.04.30.442190

**Authors:** Manish Kumar, Sandeep Kishore, David L. McLean, Yevgenia Kozorovitskiy

## Abstract

We present an open access scanned oblique plane microscopy platform *Crossbill*. It combines a new optical configuration, open hardware assembly, a systematic alignment protocol, and dedicated control software to provide a compact, versatile, high resolution single objective light-sheet microscopy platform. The demonstrated configuration yields the most affordable sub-micron resolution oblique plane microscopy system to date. We add galvanometer enabled tilt-invariant lateral scan for multi-plane, multi-Hz volumetric imaging capability. A precision translation stage extends stitched field of view to centimeter scale. The accompanying open software is optimized for *Crossbill* and can be easily extended to include alternative configurations. Using *Crossbill*, we demonstrate large volume structural fluorescence imaging with sub-micron lateral resolution in zebrafish and mouse brain sections. *Crossbill* is also capable of multiplane functional imaging, and time-lapse imaging. We suggest multiple alternative configurations to extend *Crossbill* to diverse microscopy applications.

## Introduction

Oblique plane microscopy (OPM) with its single, sample-facing objective offers unrivalled steric access in light-sheet microscopy [1]. The OPM configuration synthesizes all light-sheet microscopy associated advantages of low phototoxicity, high speed, and high resolution imaging on the foundation of the familiar epi-fluorescence microscopy configuration. It also integrates galvo scanner assisted rapid scanning in order to perform fast 3D imaging. We have shown that a plane galvo scanner, under strict physical conditions, provides a tilt-invariant scanned oblique plane illumination (SOPi), leading to geometrical distortion free 3D imaging [2]. This approach is easily extendable to two-photon microscopy [3] and is compatible with translation stage assisted hybrid scanning for large sample imaging [4]. We have also used it for deep tissue functional imaging with one-photon NIR excitation [5]. However, these advantages of OPM come at the cost of a relatively complex arrangement involving three sequentially arranged microscopy subsystems, and a scanner subsystem (**Supplementary Figure 1**). Each of these microscopy subsystems uses a high numerical aperture (NA) microscope objective. The first microscope objective (MO1) limits the maximum attainable tilt in the oblique illumination and the maximum overall system NA. The subsequent microscope objective positions (MO2 and MO3) are required to satisfy the *Herschel’s condition* and need to be positioned orthogonally in order to image the tilted intermediate image plane. This constraint leads to further reduction in the attainable system NA [1,6].

Over the last few years, multiple working OPM configurations have been demonstrated. The most common OPM configurations use an immersion objective as the first microscope objective (MO1), followed by two dry objectives as the second (MO2) and third (MO3) microscope objectives [1,3,7]. Recently, additional configurations have been implemented which enhance the overall system NA by introducing a water or solid immersion objective as MO3 [8,9]. Another configuration provides NA enhancement by placing a mirror in front of the dry MO2 to utilize the same high NA dry objective twice, as both MO2 and MO3 [10,11]. This enhanced system NA comes at the cost of unavoidable 50% loss in fluorescence signals at the polarizing beam splitter placed next to MO2. These high NA, sub-micron resolution implementations are limited to small field-of-view (FOV) imaging. Large FOV imaging, on the other hand, is made possible either by placing a diffraction grating at the intermediate image plane or by disregarding the *Herschel’s condition* and using anisotropic magnification [12,13]. Overall, a variety of configurations have resulted in a range of system NAs for the OPM family of microscopes (**Supplementary Table 1, Supplementary Figure 2**). Two observations are immediately apparent. First, there are no highly efficient configurations demonstrated between ~0.45 and ~0.95 NA; and second, all demonstrated OPM configurations rely on a dry MO2 which, when coupled with an immersion MO1, mandates the use of non-standard focal length tube or scan lens consisting of either achromatic doublets or Plossl lenses. This is problematic because high NA objectives perform best when coupled with their respective optically corrected tube lenses. Replacing standard scan lenses with Plossl lenses may limit optical performance for off-axis scan angles. The case of off-axis optical aberrations in OPM configurations is critical due to the circularly asymmetric and non-concentric overlap of MO2 and MO3 acceptance cones (**Supplementary Figure 3**). The image quality in high NA OPM configurations is adversely affected by smallest of magnification mismatches, further stressing on the requirement for standard scan and tube lenses [14]. Thus, a working OPM configuration with no compromise in the choice of tube or scan lenses is highly desirable for minimizing optical aberrations.

The OPM family of microscopes face multiple challenges for wider adoption in the research community. The absence of a commercial or open-access implementation is the primary factor keeping these microscopes out of the reach for most biology researchers. With its unconventional configuration involving three microscopy subsystems, it remains one of the most difficult microscopes to align. Addition of the galvo assisted scanning for fast 3D imaging further increases the complexity of requisite hardware and software synchronization. Finally, the high price of several parts required for the existing microscope configurations also hinders wider adaptability. Light-sheet microscopy in particular is largely driven by open access implementations and it is highly desirable to have such an implementation for the OPM family of light-sheet microscopes [15–17].

Here, we introduce a new working OPM configuration to overcome all of the challenges discussed above. We demonstrate a new arrangement consisting of three immersion microscope objectives, which enables the use of optimized optical parts offering higher performance, bridges the achievable system NA gap, and creates the most affordable OPM configuration for sub-micron resolution (**Supplementary Table 1, Supplementary Table 2**). We also provide a detailed description of the system along with assembly and alignment protocols, and open-source control software. We demonstrate the microscope’s capabilities by imaging a variety of sample types.

## Microscope design and layout

### Microscope design for cost-efficient sub-micron resolution imaging

The choice of three microscope objectives plays a major role in determining the system NA and overall cost of OPM systems (**Supplementary Table 1**). Sub-micron resolution imaging requires a microscope with greater than 0.35 effective NA. In this range, the most affordable imaging system uses the same lens as MO2 and MO3 to obtain 0.95 NA [10]. However, this approach relies on a polarizing beam splitter to redirect light leading to less than 50% fluorescence detection efficiency. In other approaches, the use of a water immersion or a solid immersion MO3 helps realize the highest system NA of 1.06 and 1.28 respectively [8,9]. However, these implementations are the most expensive, and the reliance on non-optimal tube lenses potentially adds optical aberrations and degrades achievable resolution. The *Fresnel reflection* losses in fluorescence signal from high index tilted flat surface, formed by air and the immersion media or coverslip interface in front of MO3, may become another concerning factor. One solution to this problem will require custom anti-reflection coatings on coverslips or objective surfaces designed for a specific tilt angle. However, antireflection coatings may not compensate for the reflection losses due to grazing angle incident rays; these losses are crucial to account for in estimating NA gains. For many applications very high NA imaging is not required, and a microscopy system with a moderate system NA but robust aberration and efficiency control may provide a superior choice. Our presented design is targeted towards this class of systems.

We used our open access *Crossbill Design* tool [18] to iterate through multiple configurations to arrive at the new, optimized microscope design. We observed that the constraint of non-standard tube or scan lenses arises due to different magnification and different immersion media lenses used as MO1 and MO2. This limitation can therefore be addressed by using identical MO1 and MO2 lenses. Since water immersion objectives are the most commonly used lenses for diverse biological samples, we constrain both MO1 and MO2 to water immersion objectives. The choice of MO3 with higher index immersion may appear attractive, but our design decision to reduce unwanted *Fresnel reflection* losses requires a water immersion MO3. The resulting microscope, unlike any other demonstrated designs, uses three immersion objectives. Next, we consider mechanical/steric compatibility of MO2 and MO3 and the cost factor relative to realizable effective system NA to decide on exact objective specifications. Running through all commercially available microscope objectives we conclude that 60x 1.0 NA water immersion objective is the optimal choice, offering ~0.6 effective system NA, which bridges the wide NA gap in the OPM family of microscopes. It supports a higher system NA than the most widely used OPM configuration (water immersion-dry-dry MOs), and it does so at greatly reduced cost (**Supplementary Figure 3, Supplementary Table 1, Supplementary Table 2**).

Having identified performance and cost advantages of all immersion objective configurations, we focused our attention to make every other accessory in the setup as cost-efficient as possible. For laser sources we selected single color DPSS lasers, which are larger than many compact multicolor lasers on the market, but offer efficient performance at low costs. Data acquisition card (DAQ) is important for signal generation to control and automate the microscope. We chose a python compatible, multichannel analog output USB DAQ device (USB-3101FS, MCC). The camera is one of the most important and often expensive components in any microscope. Here, we chose a high speed machine vision camera to retain cost-effectiveness (GS3-U3-23S6M-C, Flir). We wrote *Crossbill software*, which integrates the selected USB DAQ and camera seamlessly with the galvo scanners. It is equipped with an intuitive GUI to provide all required controls for operating the microscope in different imaging modes. The GUI and associated microscope hardware can all run from a laptop without the need for a powerful workstation.

### Microscope layout in a compact footprint

The OPM family of microscopes, with more than 1 meter of imaging path length between the sample and camera, occupy large space on an optical table. Here, with an optimized assembly approach, we reduce the footprint of the whole microscope to 60 cm x 60 cm x 45 cm and dramatically decrease alignment complexity. The strict requirements on distance between each optical lens pair and the galvo scanner placement makes it a challenging system for alignment. These strict distance requirements stem from the need for precise placement of the galvo scanner, with its rotation axis at the back focal plane (BFP) of MO1 and MO2, and the need for maintaining the 4f arrangement between each lens pair, in order to avoid any additional spherical aberrations in the system. We fold the microscope layout by 90° at MO1, galvo scanner, and MO2 followed by an inwards 45° fold at the MO2-MO3 interface to further save space on the optical breadboard (**Figure 1, Supplementary Figure 4, Supplementary Video 1**). The entire microscope is based on easily accessible off-the-shelf Thorlabs components (**Supplementary Table 2**). We use the smallest posts and post-holders to ensure mechanical stability. Fine adjustments, whenever required, are done with manual micro adjusters and small linear translation stages. The only custom part in this setup is a flexible water chamber between MO2 and MO3, which plays a crucial role in microscope operation and alignment (**Online Methods**). Conventional in-house microscope alignment strategies rely either on optical cage structures or on optical rails. Many researchers prefer optical cages for their four-point contact to improve stability. However, we found that neither optical cages nor rails alone are sufficient for a systematic and repeatable alignment for the OPM family of microscopes. We have developed a unique alignment protocol involving both optical cages and rails together (**Online Methods**). **Figure 1** shows the major components used in the *Crossbill* platform. We used three water immersion objectives (MO1, MO2, MO3: 60x 1.0 NA, Olympus), three scan lenses (SL1, SL2: CLS-SL, SL3: AC508-150-A-ML, Thorlabs), three tube lenses (SWTLU-C, AC508-100-A-ML, Thorlabs), two galvo scanners (GVS011, GCS001, Thorlabs), two lasers (473M50, 532M100, Dragon lasers), one CMOS camera (GS3-U3-23S6M-C, Flir), one multiband dichroic mirror (Di01-R405/488/532/635, Semrock), one dichroic mirror (FF495-Di03, Semrock), two fluorescence filters (MF525-39, MF630-69, Thorlabs), and one automated precision translation stage (MMP2, MadCityLabs). For the given choice of TL3, the camera sensor limited 2D FOV is ~338 μm x 211 μm. Galvo scanner assisted lateral scan range is limited to ~150 μm. Illumination light-sheet tilt angle in the sample is maintained at 45° and remains the same throughout the lateral scan range.

**Fig. 1.**
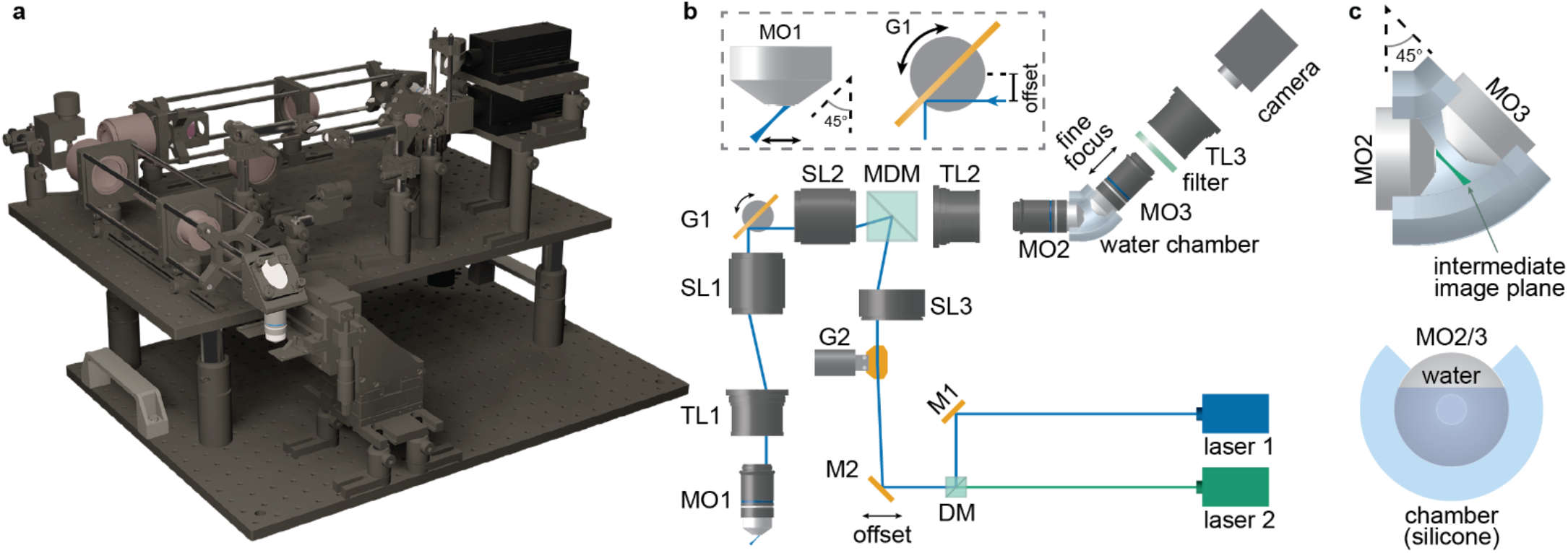
A compact single objective light-sheet microscopy configuration. A perspective 3D rendering of the assembled microscope (in **a**). A schematic showing key optical and optomechanical elements for the microscope (in **b**). The inset shows offset in the incident excitation laser beam leading to the desirable tilt in the generated light-sheet. The rotation of galvo scanner G1 is responsible for a tilt-invariant lateral scan in the light-sheet. A close-up view of the flexible/silicone water chamber to maintain water immersion for MO2 and MO3 during imaging (in **c**). MO: microscope objective, TL: tube lens, SL: scan lens, G: galvanometer scanner, MDM: multiband dichroic mirror, DM: dichroic mirror, M: mirror.

## Results

### Imaging zebrafish

Zebrafish (*Danio rerio*) are an important model organism in neuroscience, ideal for optical interrogation of neural circuit development due to their genetic tractability and transparency at early developmental stages. Compared to confocal microscopy, frequently used for high-resolution imaging of fluorescently labeled zebrafish neurons, or light-field microscopy, used for fast image collection [19], light-sheet microscopy offers significant advantages in both imaging resolution and speed [20]. *Crossbill*, with its 0.6 NA and low-cost implementation, may prove to be an efficient microscopy system for imaging embryonic and larval zebrafish. Here, we image larval zebrafish samples expressing GFP in a subset of brain and spinal cord neurons to demonstrate this capability. **Figure 2** and **Supplementary Video 2** show the imaging at 5 days post fertilization (dpf) larvae of Tg[nefma:gal4;uas:gfp] zebrafish [21]. The dataset was acquired using the *Crossbill software* at 20 fps camera frame rate. Galvo assisted lateral scan range was maintained at 100 μm. The entire zebrafish was imaged inside a 340 μm × 150 μm FOV on the camera, swept across 3400 μm length in 110 seconds. The scan switched sequentially between the galvanometer and the stage, where the galvo scanner covered 100 μm range in each sub-sweep. The acquired volumetric data were affine transformed using *transformJ* before visualization [3,22]. 3D visualization was done using ClearVolume in Fiji [23,24]. The presented visualization is based on the raw data where no post-processing, except for linear brightness and contrast adjustments, has been applied.

**Fig. 2.**
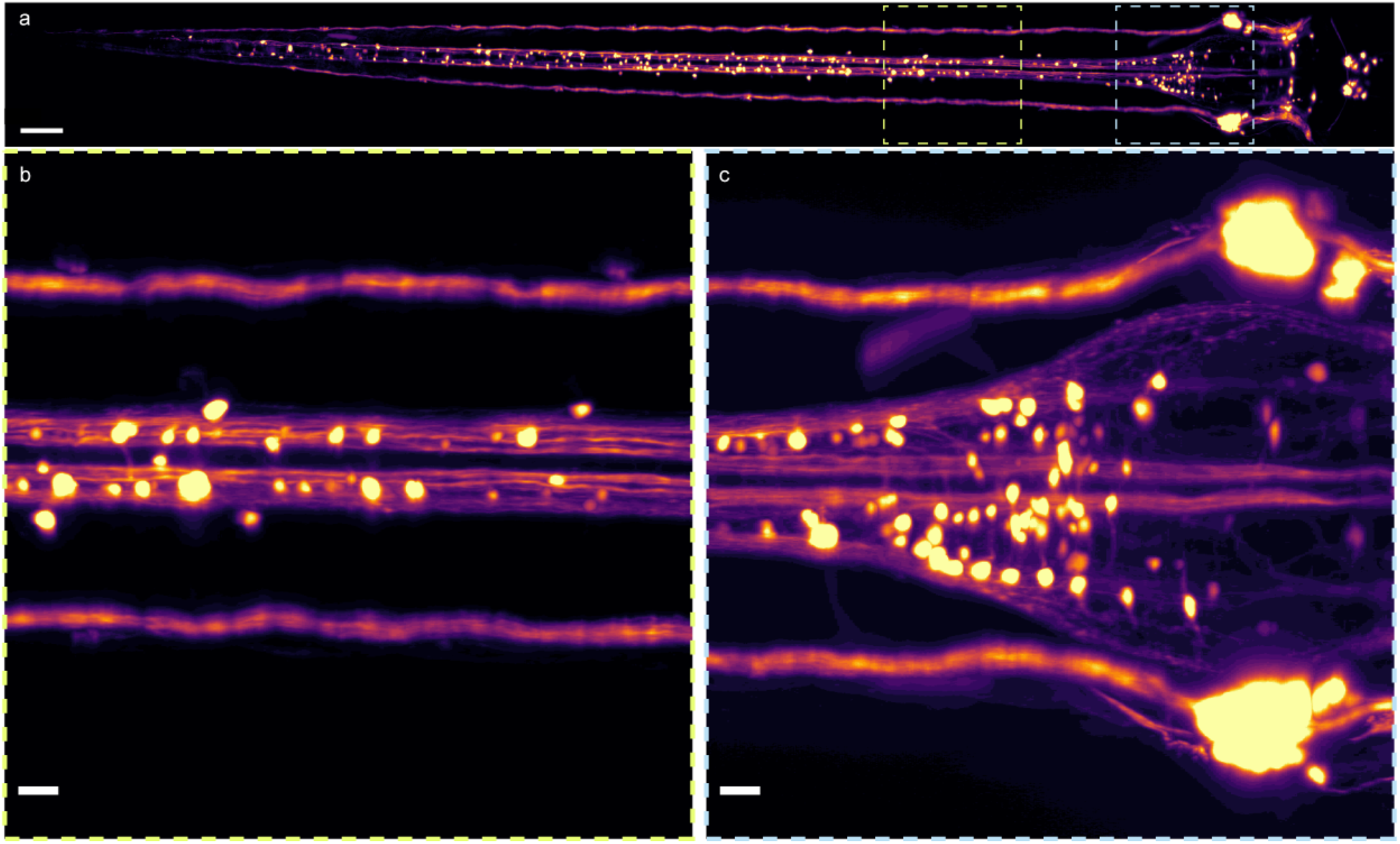
Volumetric imaging of a larval zebrafish. A 5 dpf Tg[nefma:gal4; *uas:gfp*] zebrafish larvae imaged with blue laser showing GFP expression. **(a)** A maximum intensity projection of the whole zebrafish sample imaged on *Crossbill* (scale bar: 100 μm). The projection shows a top view of the sample. **(b-c)** Enlarged view of the two highlighted regions in **a** to show the fine details (scale bar: 20 μm). **Supplementary Video 2** shows a 360° view of this dataset.

### Imaging mouse brain sections

Zebrafish larvae are relatively transparent and do not pose the challenges encountered in imaging a scattering tissue samples. Mouse brain is one example of an optically scattering sample type, broadly used across many types of neuroscience studies. One approach of imaging mouse brains with light-sheet microscopy involves post-fixation clearing steps [25]. The clearing protocols are usually slow, spanning multiple days. In the past, we have used DSLM scanning enabled single objective light-sheet microscopy to obtain cellular resolution deep tissue imaging in uncleared mouse brain samples [4]. With the current resolution upgrades, *Crossbill* platform may be useful for dendritic resolution imaging in thick mouse brain slices. As proof of principle, we imaged a *thy1gfp* mouse brain slice where a sparse subset of neurons and their fine processes is labelled with GFP. A 250 μm thick section of fixed brain tissue, containing the dentate gyrus of the hippocampus, was mounted between a coverslip and a glass slide, along with a spacer to avoid damage. This mounted slide was imaged using *Crossbill software* on our platform. We imaged a large volume region at 50 fps frame rate and 100 μm galvo assisted lateral scan range. To demonstrate the actual depth penetration and optical sectioning capability of our microscope we show imaging of densely labelled hippocampus region of the brain. **Figure 3** and **Supplementary Video 3** display this imaged region spanning 125 μm × 340 μm × 2000 μm. This entire displayed region was captured in 80 seconds and was affine transformed prior to visualizing with ClearVolume or BigDataViewer [23,26]. As for the zebrafish, the mouse brain slice data reflect only post processing with linear contrast adjustments.

**Fig. 3.**
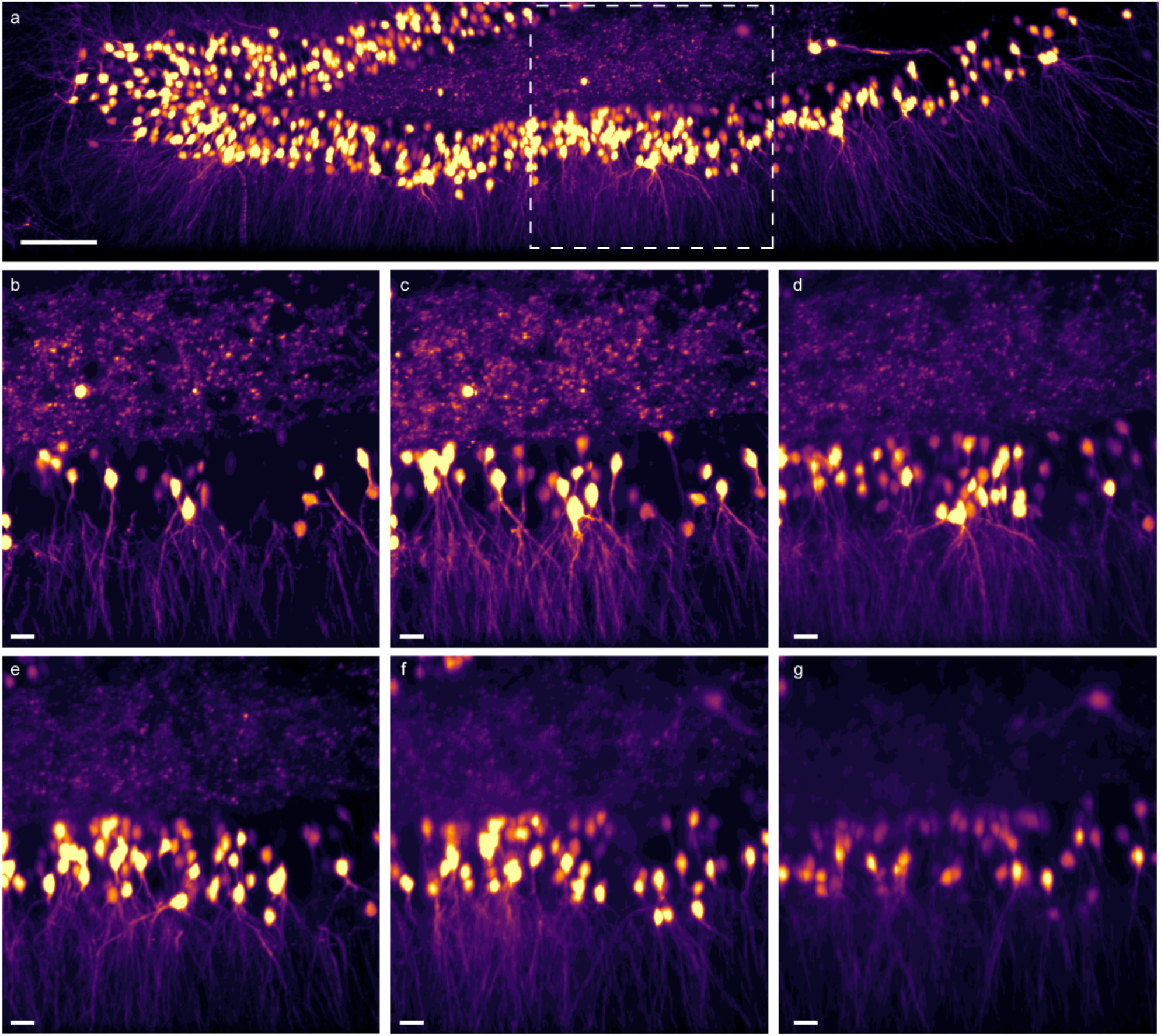
Volumetric imaging of an uncleared mouse brain section. A thy1gfp mouse brain slice (fixed, 250 μm thick) imaged with blue laser. **(a)** A maximum intensity projection of 340 μm × 2000 μm × 125 μm (width × length × height) region of the mouse brain slice showcasing neuronal cell bodies and finer projections in the dentate gyrus of the hippocampus (scale bar: 100 μm). **(b-g)** Enlarged view of the highlighted region in **a** displaying maximum intensity projection images from 0-10 μm, 10-30 μm, 30-50 μm, 50-75 μm, 75-100 μm and 100-125 μm depth respectively (scale bar: 20 μm). **Supplementary Video 3** shows a 360° view of this dataset.

## Discussion

We present *Crossbill* as the most affordable and compact single objective light-sheet microscopy platform for sub-micron resolution imaging. It attains a moderately high effective system NA of 0.6 which fills the wide gap in system NA for photon-efficient implementations in this class of microscopes to date. This small footprint microscope, with its ability to perform large scale structural, rapid multi-plane functional, and long term timelapse imaging, is a versatile imaging platform. The *Crossbill* platform comes with open access design, assembly and operation tools, making it easy to replicate, adapt, and upgrade.

Unlike conventional optical instruments, SOPi based microscopes cannot be aligned with standard procedures. A non-scanned single objective light-sheet does not pose the same alignment challenges. We have previously demonstrated the importance of precise placement of the galvanometer mirror in the setup. The rotation axis must match the back-focal plane of the microscope objective (MO1 and MO2) [2]. This is essential for tilt-invariant lateral scanning of the light-sheet. This condition, combined with the need to ensure minimum optical aberrations in the system, requires a strict 4f arrangement between each optical lens pair. In the past, attempts to adapt an existing microscope body to form the microscope have been made [8], but in a commercial microscope body the objective-tube lens pair does not form a 4f configuration, making it a sub-optimal approach. Among the many custom designs, optical rails or cage structures are popular for alignment assistance. Some light-sheet microscope designs use optical rails for alignment [15], and most rely on cage structures [3,4,8,9,25,27]. While the existing alignment strategies work, they are insufficient for systematic and easy alignment for the galvo-assisted lateral scanning OPM family of microscopes. Over the past years, we have refined and simplified our alignment strategy to make it systematic and reliable. Unlike other alignment strategies that use either cage structures or optical rails, our alignment strategy relies on both. The alignment strategy also integrates *Crossbill software* for facilitating fine alignment. We have ensured that the alignment protocol does not rely on expensive alignment tools, using inexpensive custom tools without quality compromises. The alignment protocol has been validated in the absence of an optical table, illustrating robustness of the procedures. The success of this alignment protocol enabled folding the optical path of the microscope into a compact footprint. In the future, the footprint can be further reduced by adding folds along the third axis and selecting a compact multicolor laser system.

There are multiple approaches to generate a light-sheet. Here, we rely on a DSLM, which reduces shadow artifacts and allows superior contrast for deeper imaging in light-scattering samples [4,28]. DSLM approach with no slit integration supports the use of all of the available laser power for excitation. In the present setup we did not vary laser beam size to change illumination NA. If required, it can be accomplished by introducing a beam expander combined with a variable slit aperture. The integration of a rolling shutter enabled sCMOS camera would help with synchronized confocal slit detection, to further enhance the resolution and optical sectioning capabilities of the microscope [29]. The use of an sCMOS camera would also enhance the dynamic range and sensitivity, leading to faster imaging. As common in other light-sheet microscopy approaches, deconvolution with a measured point-spread-function can be used to further enhance the attainable resolution and contrast [8,9,11,27].

The *Crossbill* platform is not limited to the single design presented here but is built with scalability in mind. Other existing or new designs can be readily implemented. The combined use of cages and optical rails is extendable to any design which requires precise placement of optical elements with respect to a scanner. The *Crossbill software* has the same flexibility as *Crossbill design*, where a new configuration file could be entered by a user to make the software control a new microscope design. We believe the present example implementation demonstrates an optimal performance to cost ratio, but the tools provided equip a reader to improve the microscope design as per application demands and build alternative versions of the microscope. We provide a list of several alternate designs (**Supplementary Note 3**) to make the OPM family of microscopes accessible for the broader biology research community.

## Acknowledgements

We thank Lindsey Butler for mouse colony management, and Deanna Badong and Vasin Dumrongprechachan for assistance with sample preparation.

## Online Methods

### *Crossbill* assembly

A two-layer aluminium breadboard assembly (MB4560, Thorlabs) held together with 1” posts and post-holders (RSHT4, RS150, Thorlabs) form the base structure for *Crossbill*. Three dovetail optical rails (RLA300/RLA150, Thorlabs) are fixed on the upper breadboard to mount three parts of the imaging arm of the *Crossbill*. A galvo scanner (GVS211) is mounted at the common corner of the first two optical rails. The first part of the imaging arm consisting of MO1 (LUMPLFLN60XW, Olympus), tube lens TL1 (SWTLU-C, Olympus), and scan lens SL1 (CLS-SL, Thorlabs) is mounted on the first optical rail with the help of standard cage structures and optical posts (LCP08, LCP02, KCB1EC, ER12, ER3, PH20, TR20, RC1, Thorlabs). The second part of the imaging arm consisting of the same optomechanical elements as the first part is mounted on the second optical rail. The third part of the imaging arm consisting of MO3 (LUMPLFLN60XW, Olympus), tube lens (AC508-100-A-ML, Thorlabs), camera (GS3-U3-23S6M-C, Flir), and fluorescence filters (MF525-39, MF620-52, Thorlabs) is mounted on the third optical rail with the help of cage structure mounts and optical posts (LCP02, LCP08, ER6, ER1.5, CFS1M, PH20, TR20, RC1, Thorlabs). MO3 is mounted on a cage compatible z-translation mount (SM1Z, Thorlabs) to enable fine focusing on the intermediate image plane. For the illumination arm, two lasers (MGL-III-532-100mW, MBL-III-473-50mW, DragonLasers) are vertically aligned on two additional levels on the upper aluminium breadboard with the help of small breadboards and optical posts (MB1015, PH30, PH100, TR30, TR100, Thorlabs). These lasers are combined together and coaligned with a dichroic mirror (FF495-Di03, Semrock) and a mirror mounted on optical posts (ME1-G01, C45P, ER3, CM1-DCH, CP02F, TR100, PH100, Thorlabs). The combined beams are then directed through a pair of mirrors to a galvo scanner (GVS201, Thorlabs) which redirects the beam towards scan lens SL3 (AC-508-150-A-ML, Thorlabs). The two mirrors, prior to the galvo scanner in the laser path, are mounted on a manual translation stage to help control the offset in the laser beam. Beyond the SL3 lens, the laser beams are redirected into the main imaging arm with a multi-band dichroic mirror (Di01-405-488-532-635, Semrock) inside a cage compatible mount (CM1-DCH, Thorlabs) placed next to the second scan lens SL2. A precision 2-axis motorized translation stage with the third axis as manual adjustment (MMP3_manualZ, MadCityLabs) is placed on the lower breadboard to access the image volume of MO1. Two handles are fixed on the lower breadboard for easy transportation of the assembled setup. The whole *Crossbill* assembly is shown in **Supplementary Video 1**. To control and synchronize the galvo scanners and the camera, a multichannel analog output USB DAQ card (USB-3101FS, Measurement Computing) is used. **Supplementary Table 2** lists all major components, along with current price estimates.

### Flexible water chamber

A water chamber is required between MO2 and MO3 to contain the immersion media during imaging. This water chamber is cast into a flexible silicone material to assist with easy alignment of the microscope. A 3D model of the chamber was prepared using Blender (blender.org, Blender Foundation). The chamber has a ‘C’ shaped cross-section and is modeled to accommodate MO2 and MO3 objectives at a 45° tilt. A negative mold of the chamber was created in Blender with thin walls and a top opening. This negative mold is 3D printed with PLA material and the resulting physical mold is filled with a silicone elastomer (Sylgard 184, Dow) and cured at room temperature for 48 hours. Once cured, the outer PLA mold is removed with a knife to recover the silicone water chamber. This chamber is secured in the setup with the help of two 3D printed PLA clips and fitting cap screws with nuts.

### Light-sheet span and tilt management

A digitally scanned light-sheet requires an additional galvo scanner in the setup. Cylindrical or Powell lenses could also be used to generate the light-sheet. However, the galvo assisted approach supports digital control of the light-sheet span, via settings available on the *Crossbill software* GUI. The addition of manual translation stage mounted mirrors enables control over the offset in the incident beam, allowing light-sheet tilt management. This feature supports easy switching between the on-axis alignment beam and the off-axis imaging beam arrangements.

### *Crossbill* alignment

The alignment steps are split into three parts. The first part deals with the alignment of all elements between MO1 and MO2, the second part consists of aligning optical elements between the MO3 and the camera, and the final part relates to the illumination path alignment. For the **first part**, all lenses are removed from the setup leaving behind the corner mirrors (**Supplementary Figure 6a**), including the galvo scanner mirror G1. A visible laser beam (Green, 532M100, Dragon Lasers) is passed through the setup from the MO1 end to exit at the MO2 end. Cage alignment plates (CPA1, LCPA1, Thorlabs) are used to ensure that the incident laser beam is centered in the MO1-SL1 arm. The position of the first galvo G1 is adjusted to make the laser beam reflect from the center of G1 mirror and pass through the center in the SL2-MO2 arm. The centering of the laser beam is ensured with cage alignment plates. Position of the centered laser beam is marked on a screen kept >0.5 meters away from the MO2 end. Next, the laser is expanded into a collimated beam with a microscope objective (LUMPLFLN60XW, Olympus) and an achromatic doublet (AC508-100-A-ML, Thorlabs). This collimation is confirmed with a custom made *trishanku* (or crux) collimation tester [30]. The first tube lens TL1 and scan lens SL1 are inserted in the setup and the collimated beam is made to pass through them. The distance between TL1 and SL1 is fine-tuned to retain a perfectly collimated beam coming out the other end (**Supplementary Figure 6b**). Once confirmed with *trishanku*, this distance between TL1-SL1 lens pair is locked by tightening the screws in the corresponding cage plates on the cage structure rods. The temporary beam expander arrangement is removed to recover the un-expanded laser beam. Next, a **laser-in-laser-out** alignment strategy is performed to ensure all the optical elements between MO1 and MO2 are at perfect distance, where each of the lens pairs forms a 4f setup. In brief, the **laser-in-laser-out** strategy is based on a low divergence laser beam entering the system at MO1 end that is expected to emerge as a low divergence laser beam at the MO2 end. For this, MO1 is inserted and its position is fine-tuned to obtain a perfectly collimated beam beyond TL1 (**Supplementary Figure 6c**). The collimation is verified with a temporary plane mirror inserted at approximately 45° between TL1 and SL1 to reflect the beam and using *trishanku* in the reflected beam path. Once confirmed, MO1 is locked into this position by tightening the screws in the corresponding cage plates. This completes the alignment of the MO1-TL1-SL1 alignment in the setup.

Next, SL2 is inserted in place (**Supplementary Figure 6d**), the temporary plane mirror is moved to a new location beyond SL2, and SL2 is fine adjusted to obtain a collimated beam beyond it (tested and confirmed with *trishanku*). Once collimated, the temporary plane mirror is removed and TL2 is inserted (**Supplementary Figure 6e**). To fine-tune TL2 position, first SL1 is carefully removed with its cage plate steady, marking its position, and then TL2 is repositioned to obtain a collimated beam. The collimation was confirmed with *trishanku* placed beyond TL2. SL1 is carefully re-inserted in its marked previous position and now MO2 is inserted in the setup (**Supplementary Figure 6f**). A diverging laser beam indicates the error in MO2 position and it can be corrected by fine adjusting MO2 to obtain non-diverging laser beam output. Once a non-diverging beam output is obtained, each of the elements in the SL2-MO2 arm should be confirmed to be locked in its respective position. This beam matches the pre-marked position on the distant screen to confirm that all the elements are center-aligned. This completes the **laser-in-laser-out** strategy to ensure the optimal distance between each optical lens pair between MO1 and MO2, along with the objectives.

The next step is to ensure that the galvo scanner G1 is placed in the optimal position to obtain tilt-invariant scanning. For this, the first galvo scanner is driven by a low frequency (~1 Hz) ramp signal, generated through the *Crossbill software*, and the laser beam coming out of MO2 is observed on a distant screen. The beam moves along horizontally for the incorrect placement of the galvo scanner. At this point, the advantage of using both rails and cage structures is evident. While cage structure help keep the optical elements locked at a fixed distance, the optical rails allow for the movement of MO1-SL1 and SL2-MO2 segments with respect to the galvo scanner G1 (**Supplementary Figure 6f**). Laser beam divergence indicates incorrect distance between MO1-SL1 and SL2-MO2 segments. Rail carrier positioners (RCN, RCN1, Thorlabs) are used for precise micromovement of the MO1-SL1 and SL2-MO2 segments. The position corresponding to a constant laser beam displacement on the screen indicates the optimal relative positions of the MO1-SL1 and SL2-MO2 segments. This laser beam displacement, typically less than a millimeter and hence largely undetectable by eye, remains constant irrespective of the screen distance. This characteristic follows from the tilt-invariant scan principle of the SOPi microscopy, which has a simple geometrical proof [2]. This point marks the completion of the first part of the alignment process.

The **second part** deals with MO3-camera alignment. The camera is removed, and the rest of the assembly, except for the optical rail, is moved to the bottom breadboard with the MO3 end facing the unexpanded alignment laser beam. The MO3-TL3 distance is adjusted to ensure an expanded collimated beam output from the TL3 end. MO3 is removed, keeping its exact position marked through the z-axis translation mount attached to the cage structure. The camera is connected to a computer to obtain live feed and, while keeping TL3 fixed in its place, the camera position is adjusted to sharply image a distant object visible outside window. Alternatively, when outdoor view is inaccessible, an expanded and collimated laser beam is used with TL3-camera arrangement, and the camera is fine positioned to form a sharp point focus on the camera sensor plane. Here, the laser power is reduced with a neutral density filter for an unsaturated view of the focused spot on the camera sensor. Once adjusted, the camera is fixed in this position and MO3 is inserted back. This MO3-camera assembly is then fixed on top of the third optical rail in the setup. The diagonal holes on the upper breadboard are used as a guide for a precise 45° orientation of the MO3-camera assembly. The flexible water camber is attached to bring MO2 and MO3 together and the chamber is filled with water to immerse both the microscope objective lenses. A cage alignment plate (LCPA1, Thorlabs) is inserted between SL2 and TL2, roughly matching the common focal plane of both lenses. MO3-camera assembly is moved to bring the image of the hole of the alignment plate into the center of the view of the camera. This step requires an in-plane movement of the assembly, where the optical rail orientation is maintained at 45° (**Supplementary Figure 6g**).

The **third part** of the alignment process positions the illumination laser beams. The cage alignment plate between SL2 and TL2 lenses (used in the previous part) is removed and a multiband dichroic mirror is inserted adjacent to the SL2 lens. SL1 is removed and SL3 lens is introduced, with its position adjusted to collimate the dichroic reflected beam (**Supplementary Figure 6h**). Collimation is confirmed with *trishanku*. SL1 is re-inserted in its position to make the laser beam converge beyond SL3. The second galvo scanner G2 is placed with its rotation axis centered at the laser beam convergent point (**Supplementary Figure 6i**). The galvo reflected beam is redirected towards the imaging laser(s) with the help of two orthogonally arranged, translation stage mounted mirrors (shown as a single mirror M2 for the simplicity in the schematic diagram **Figure 1**). MO1 is removed to allow the unexpanded alignment laser beam to counter-propagate towards the imaging lasers. Both the imaging lasers are turned on and merged together with a mirror and a dichroic mirror, to co-align with the alignment laser beam path. MO1 is re-inserted in its fixed position and the alignment laser is turned off. Imaging laser beams follow the on-axis path of each optical lens to exit MO1 along the axial direction. M2 mirror is offset with the help of a translation stage to create the 45° oblique beam in front of MO1 (**Supplementary Figure 6j,k**).

The next step consists of ensuring that MO3 working distance is matched to the intermediate image plane. An agar embedded microbead sample is placed in front of MO1. The sample surface is brought closer to MO1 with a few water drops to ensure water immersion based optimal imaging conditions. Camera live feed is observed to obtain the images of fluorescent microbeads. Out of focus beads images indicate mismatched positioning of MO3 with respect to the intermediate image plane. MO3 is adjusted along its axis with the help of the corresponding cage compatible z-translation mount until the image of microbeads comes into sharp focus. The tilt-invariant scanning of the light-sheet is confirmed with a standard clean coverslip imaged without a fluorescence filter, as detailed elsewhere [2].

### Crossbill software

A dedicated software was written in python, an open-source programming language. This is a deviation from the traditional approaches in the scientific community, where control software is usually written in LabVIEW or MATLAB. Many projects use the micromanager platform [31] for its camera and translation stage compatibility. However, SOPi microscopy requires high levels of synchronization between the galvo scanner position and the corresponding camera acquisition trigger. There are many parameters which directly control the 3D FOV and imaging speed of the microscope, and they need to be controlled on-the-fly with user interaction. Thus, we created a complete software with a dedicated GUI to perform these specific tasks. The software directly connects with the camera, DAQ card, and translation stage. It provides suggested imaging modes for structural, functional, and time-lapse imaging. It is written with a multithreading option to enable optimized parallel performance on a low-end laptop. The software can also operate with a camera connected to micromanager, or other programs, where *Crossbill software* simply controls frame acquisition through DAQ generated camera triggers.

### Zebrafish larvae preparation

All procedures for zebrafish imaging conform to NIH guidelines on animal experimentation and were approved by the Northwestern University Institutional Animal Care and Use Committee. Experiments were performed using 5-day-old transgenic zebrafish larvae, Tg[nefma:gal4;uas:GFP] [21], when they have a fully inflated swim bladder and are free-swimming. Briefly, larvae were anesthetized in 0.02% w/v ethyl 3-amino-benzoate methanesulfonic acid (MS-222; Sigma Aldrich), transferred to a glass-bottomed dish, and embedded upright in low-melting-point agar (1.4% in system water). After the agar had solidified, it was layered with more anesthetic solution to prevent agar desiccation and ensure that fish remained anesthetized and stationary for imaging experiments.

### Mouse brain slice preparation

Animals were handled according to protocols approved by the Northwestern University Animal Care and Use Committee. All mice were group-housed, with standard feeding, light-dark cycle, and enrichment procedures. Young adult Thy1-EGFP M line mice (Jax, 007788) were used in the study. Mice were deeply anaesthetized with isoflurane and transcardially perfused with 4% paraformaldehyde (PFA) in 0.1 M phosphate buffered saline (PBS). Brains were post-fixed for 1 day and washed in PBS, prior to sectioning at 250 μm on a vibratome (Leica Biosystems), mounting, and coverslipping using Glycerol:TBS (3:1) with a spacer.

### Stitching and visualization

There was no need to perform stitching for the presented data. The *Crossbill software* controls the motion of the galvo assisted scan and the translation stage to provide already stitched data. The only step required prior to the visualization was an *affine transformation* [22]. The data were visualized with ClearVolume or BigDataViewer in Fiji/imageJ [23,24,26,32].

## Supplementary Information

**Supplementary Figure 1.**
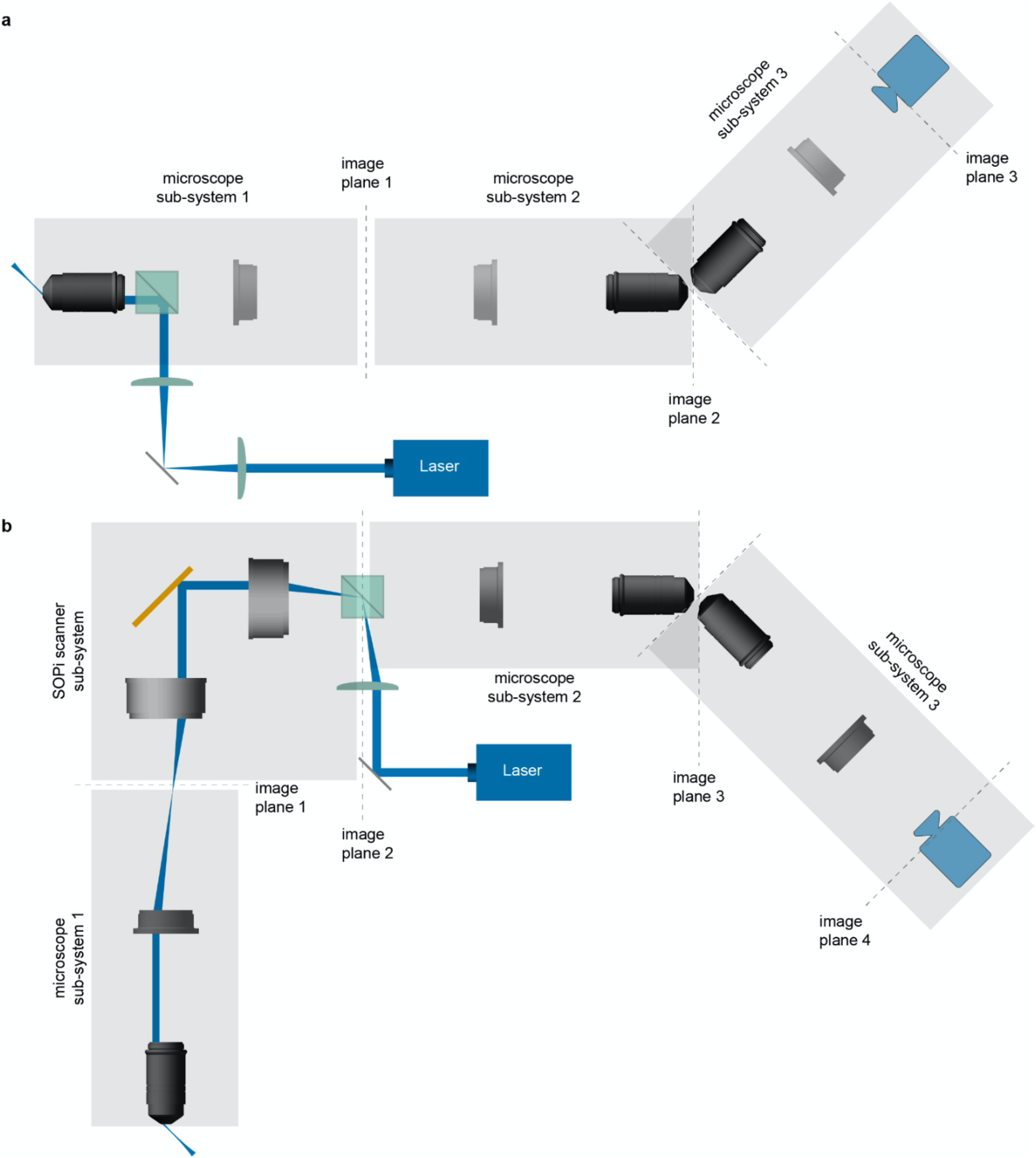
The optical arrangement of a single objective light-sheet microscope. **(a)** The OPM configuration consists of three microscopy sub-systems. First two sub-systems together satisfy both *Herschel’s* and *sine* conditions to form an intermediate image of an optically sectioned plane. The third microscopy sub-system magnifies and images this intermediate image plane on a camera sensor. **(b)** The SOPi configuration introduces a 4f scanner subsystem, consisting of two scan lenses and a galvanometer scanner in the OPM configuration, to enable tilt-invariant lateral scan of the generated light-sheet.

**Supplementary Table 1.** Existing objective combinations for the OPM family of microscopes. The table lists currently demonstrated configurations in the ascending order of system NA. While both dry and immersion objective choices have been demonstrated for MO1 and MO3, no configuration has ever been demonstrated with an immersion MO2.

| Author(s), Journal, Year | MO1 | MO2 | MO3 | NA<sub>eff</sub> |
| --- | --- | --- | --- | --- |
|  | Dry | Dry | Dry |  |
| Hoffmann et al., Optica, 2019 | 4x/0.28 | 4x/0.28 | 4x/0.28 | 0.28 |
|  | Oil/Water | Dry | Dry |  |
| Bouchard et al., Nat. Photonics, 2015 | 20x/1.0 | 20x/0.75 | 20x/0.4 | 0.33 |
| Kumar et al., Opt. Express, 2018 | 20x/1.0 | 20x/0.75 | 20x/0.45 | 0.35 |
| Dunsby, Opt. Express, 2008 | 60x/1.35 | 40x/0.85 | 10x/0.3 | 0.45 |
| Sparks et al., Biomed Opt Express, 2020 | 40x/1.15 | 20x/0.75 | 20x/0.75 | 0.58 |
| Kim et al., Nat. Methods, 2019 | 60x/1.20 | 50x/0.95 | 50x/0.95 | 0.95 |
|  | Water | Dry | Water |  |
| Yang et al., Nat. Methods, 2019 | 60x/1.27 | 100x/0.9 | 60x/1.0 | 1.06 |
|  | Silicone | Dry | Glass |  |
| Sapoznik et al., eLife, 2020 | 100x/1.35 | 40x/0.95 | 20x/1.0 | 1.28 |

**Supplementary Figure 2.**
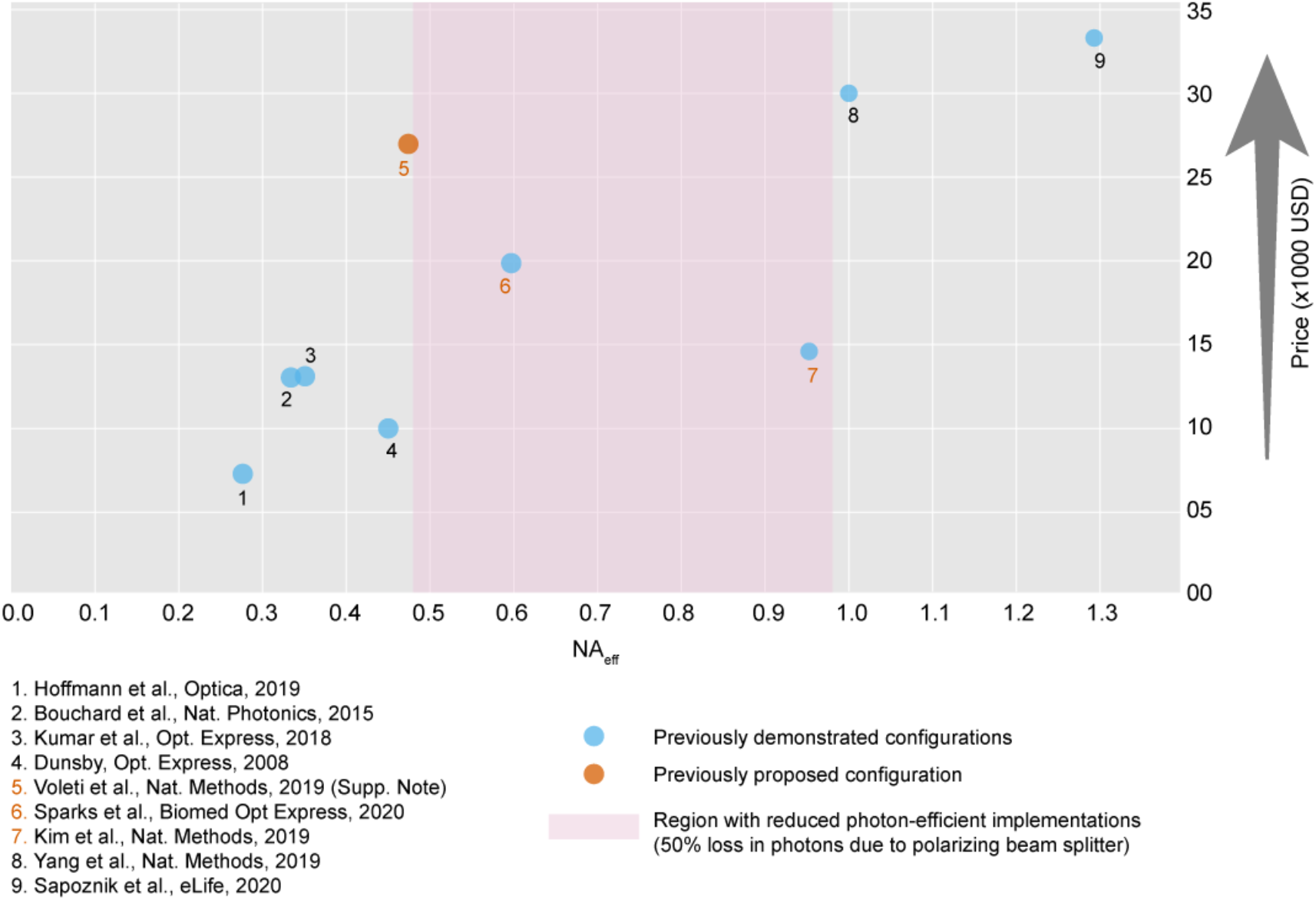
Effective system NA and price comparison among existing single objective lightsheet configurations. The microscope objectives used in the configurations are listed in **Supplementary Table 1** and further information is available in the respective publications [1,3,7–12,27]. The red region between 0.45 and 1.0 highlights unavailability of a photon-efficient OPM configuration. Price estimates include three microscope objectives without other accessories.

**Supplementary Table 2.** Crossbill parts list along with current price estimates.

| Item | Part Number | Qty | Price |
| --- | --- | --- | --- |
| Microscope objectives | LUMPLFLN60XW, Olympus | 3 | ~\$9,600 |
| Tube lens | SWTLU-C, Olympus | 2 | \$800 |
| Scan lens | CLS-SL, Thorlabs | 2 | \$5,600 |
| Large beam galvo | GVS011, Thorlabs | 1 | \$1,600 |
| Small beam galvo | GVS001, Thorlabs | 1 | \$1,050 |
| Achromatic lens | AC508-150-A-ML, Thorlabs | 1 | \$150 |
| Achromatic lens | AC508-100-A-ML, Thorlabs | 1 | \$150 |
| Blue laser | 473M50, Dragon lasers | 1 | \$1,160 |
| Green laser | 532M100, Dragon lasers | 1 | \$980 |
| Dichroic mirror | Di01-R405/488/532/635, Semrock | 1 | \$615 |
| Dichroic mirror | FF495-Di03, Semrock | 1 | \$285 |
| Fluorescence filter | MF525-39, Thorlabs | 1 | \$260 |
| Fluorescence filter | MF630-69, Thorlabs | 1 | \$260 |
| Optical breadboard | MB4560, Thorlabs | 2 | \$800 |
| Z-Axis Translation Mount | SM1Z, Thorlabs | 1 | \$210 |
| Miscellaneous optomechanical mounts | Thorlabs | - | \$2,000 |
| DAQ board | USB-3101FS, MCC | 1 | \$550 |
| Camera | GS3-U3-23S6M-C, Flir | 1 | \$1,050 |
| **Total** | | | **\$27,120** |
| Manual translation stage (option 1) | LT3, Thorlabs | 1 | \$1,350 |
| Automated translation stage (option 2) | MMP2 (custom.), MadCityLabs | 1 | ~\$8,000 |
| Laptop/Desktop | 64 bit Windows 10, 8+ GB RAM, 500+ GB storage | 1 | ~\$1,000 |
| **Max. total** | | | **\$36,120** |

**Supplementary Figure 3.**
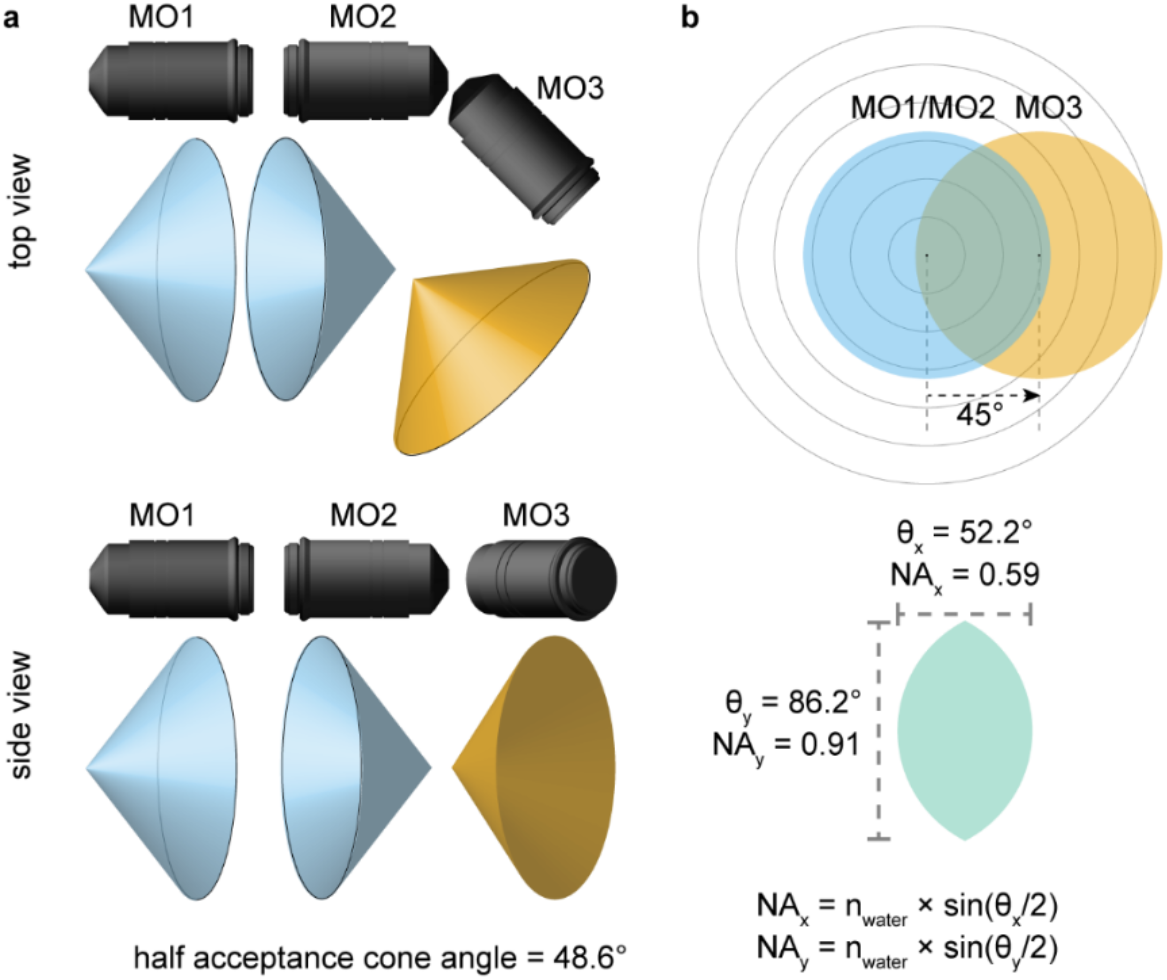
Effective system NA for the presented configuration. **(a)** Top and side views of three water immersion microscope objectives of 1.0 NA used in the current configuration. The acceptance cones of the microscope objectives are also shown. **(b)** Calculation of the effective system NA consists of the tilt angle between MO2 and MO3 leading to a reduction in the system NA. As expected, the reduction in NA is larger along the tilt affected direction.

**Supplementary Figure 4.**
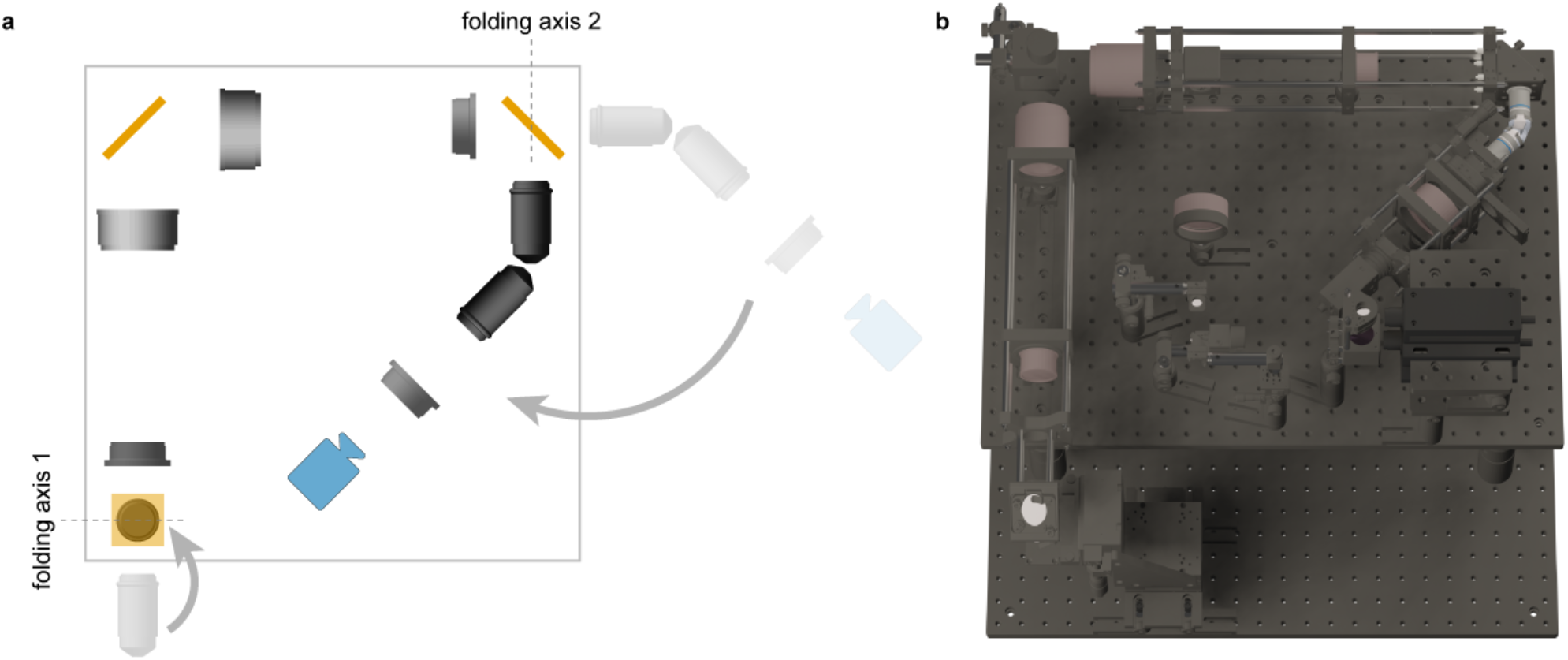
Folding SOPi configuration to a compact format. **(a)** Schematics showing how two additional folds in the SOPi configuration help reduce the overall footprint of the system. **(b)** A top view (computer generated 3D rendering) of the assembled compact Crossbill system.

**Supplementary Figure 5.**
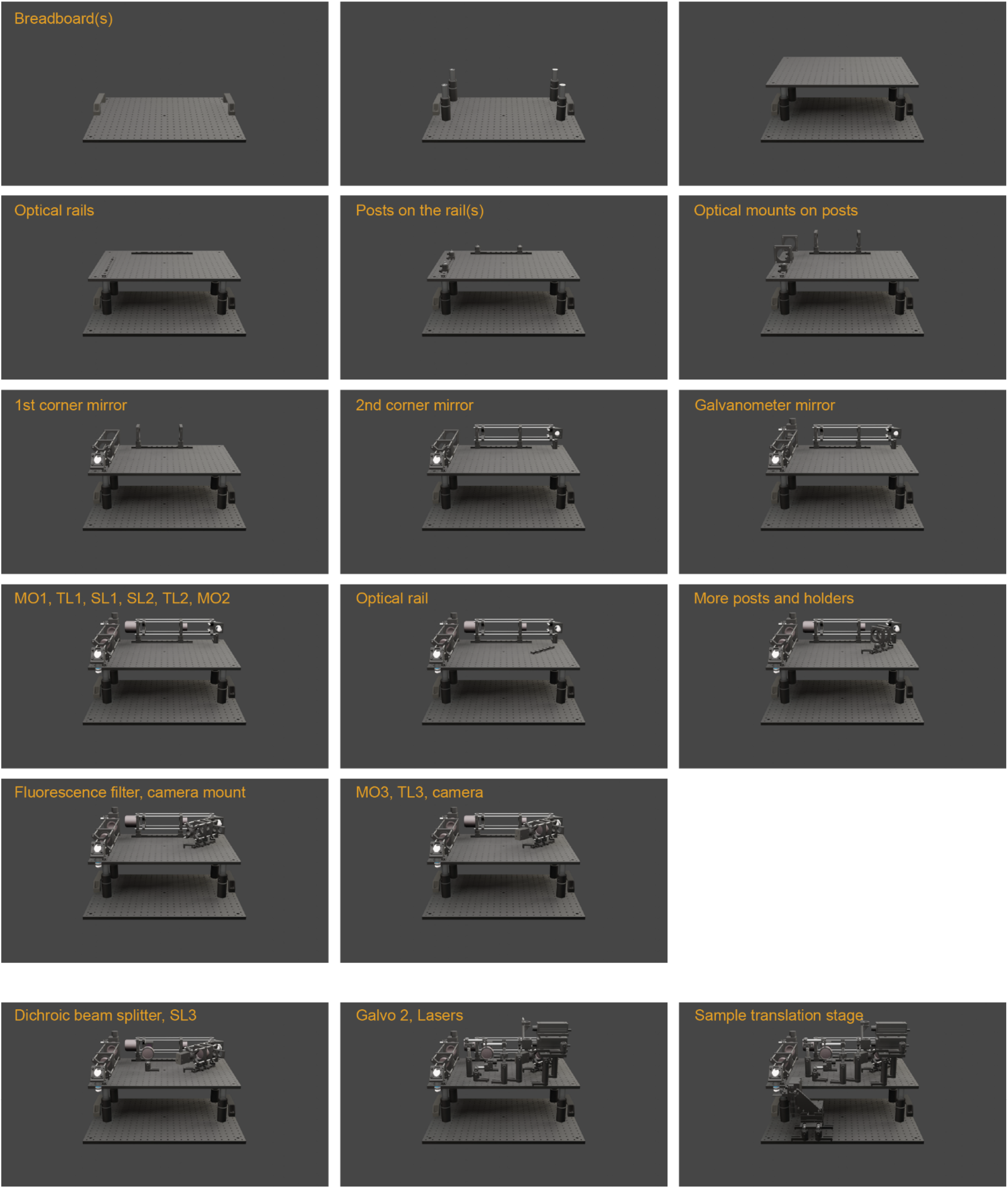
Assembly steps for *Crossbill* platform. The whole platform fits in 60 cm × 60 cm × 45 cm space. Associated parts list is given in the **Supplementary Table 2.**

**Supplementary Figure 6.**
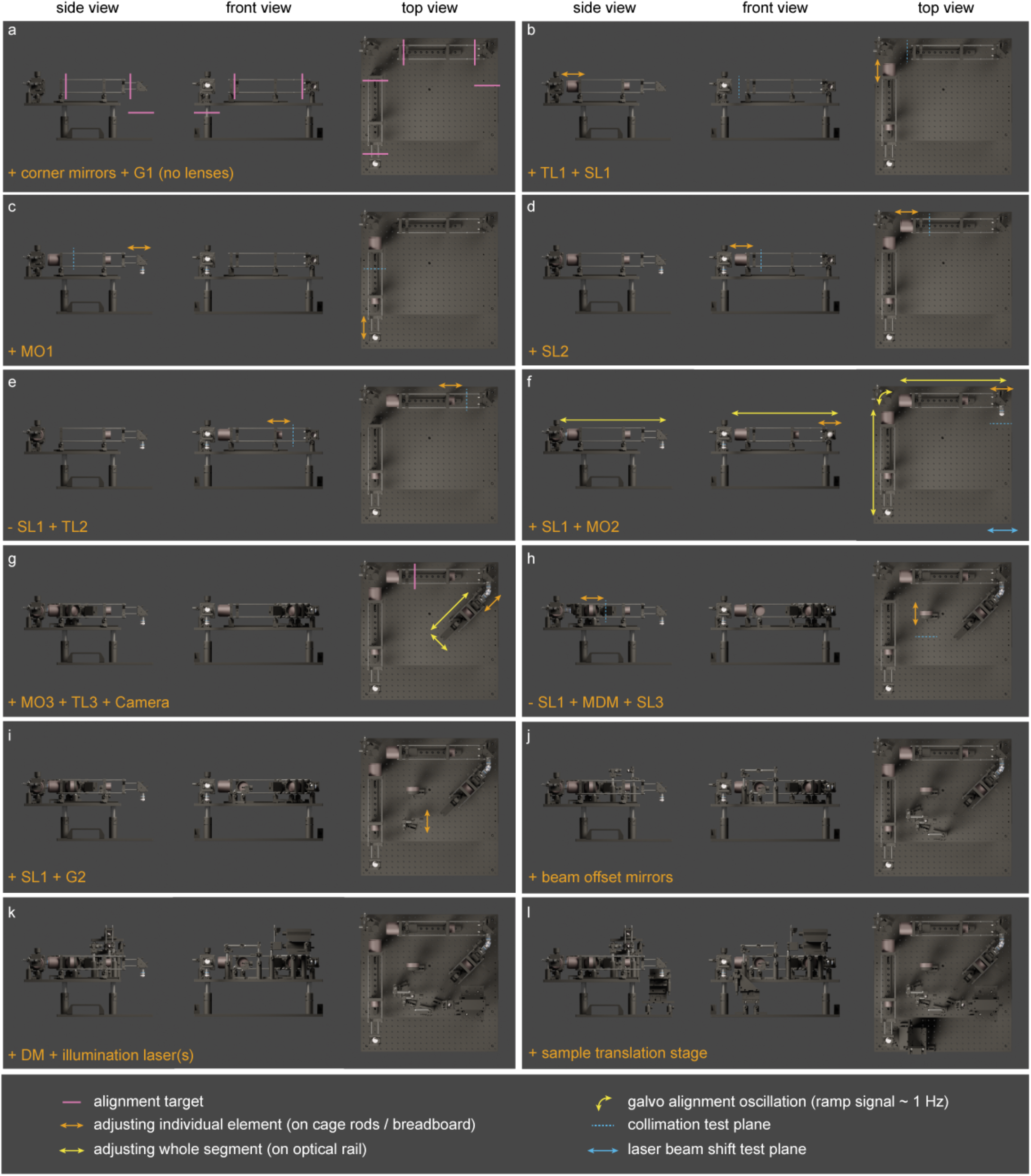
Alignment steps for *Crossbill* platform.

**Supplementary Note 1 | Design steps with *Crossbill Design***

*Crossbill Design* is a user-friendly GUI to quickly design OPM configurations [18]. **Supplementary Figure 7** shows a screenshot of the current design used in *Crossbill*. The choice of optical elements in **Supplementary Table 2** are as per design parameters displayed here. *Crossbill Design* also provides information about effective magnification, effective NA, and maximum field of view. For the design presented in this work we have the following:

- Effective system magnification: 33.33x
- Effective NA: 0.59 (for NA_x_). The current implementation of *Crossbill Design* does not estimate effective NAy precisely. This is sufficient because NAy, being larger than NA_x_, does not limit the effective system NA. An accurate NAy is calculated in **Supplementary Figure 3**.
- Beam offset: 2 mm. This is the amount of lateral beam-offset required to obtain the desirable tilt (45° in our case) in the light-sheet. The offset needs to get introduced prior to the scan-lens SL3.
- Field of view: 442 μm (maximum). This is the maximum field of view in the lateral plane. The observed field of view is largely dependent on the lateral scan range and other parameters [cite]. In our demonstrated design we have prioritized greater sampling, resulting in a 338 μm × 211 μm field of view.

**Supplementary Figure 7.**
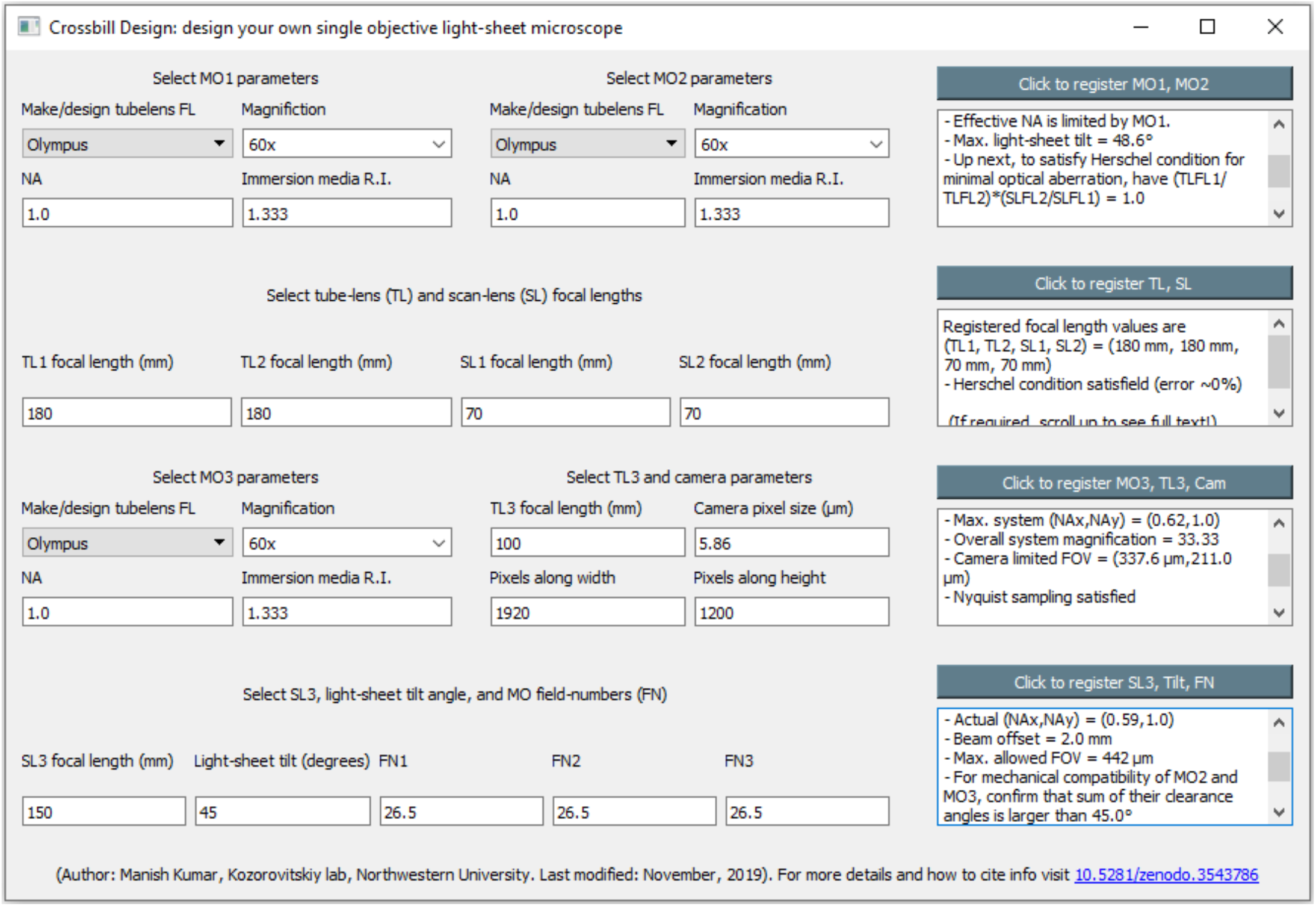
*Crossbill Design* tool for generating the new working configuration presented here.

**Supplementary Note 2 | *Crossbill software***

The software consists of modules for each hardware component and multiple readable functions for various methods. The OS support and python compatibility depends on the respective manufacturers of the chosen hardware. If the hardware is supported, a user can easily overwrite the relevant lines, as in our openly available code, to the device operational in their microscope. We will continue expanding the list of supported python-compatible hardware. The *Crossbill software* offers the ease of use comparable to a commercial system but differs in that the source code is readable, openly available, and modifiable to adapt to a different set of hardware. We believe that *Crossbill* will encourage microscopy builders to not only innovate with hardware but also create new, feature packed open software.

In principle, micromanager [31] (or pycro-manager [33] for python support) can also be used for control and automation of each of the hardware in our microscope. However, it is not simple to generate on-demand DAQ signals of desirable shape, frequency, and amplitude with micromanager. Given the complexity and precision requirements of control signals for the camera and galvo scanners, and their direct implications for imaging parameters, it is essential to have a dedicated software interface streamlined for this task. A future integration of pycro-manager with *Crossbill* will help expand the list of compatible devices. **Supplementary Video 4** shows *Crossbill software* in action during a simple image acquisition.

**Supplementary Note 3 | Alternate designs/upgrades**

The configuration demonstrated in this work, being focused on a low-cost implementation, makes seceral considered compromises. Many alternate configurations and upgrades, driven by a particular application demand, can be implemented using the same *Crossbill* platform. Using *Crossbill Design*, it is possible to quickly cycle through various combinations of microscope objectives from different vendors. Microscope objectives from Nikon, Mitutoyo, and a few others, require a longer 200 mm focal length tube lens and therefore result in an increase in the microscope footprint. A slightly larger off-the-shelf (MB4575/MB6060, Thorlabs) or custom breadboard can be used in such cases. Since our platform uses optical rails for housing components, it is possible to mill a custom platform with bare minimum number of tapped holes to mount optical rails, where the rest of the optics fit on top of the optical rails. Wider optical rails (34 mm/ 66 mm/ 95 mm, Thorlabs) may be used for better mechanical stability and smoother motion of the optical components.

### Alternate cameras

The current configuration uses a machine vision camera which is limited in its sensitivity and imaging speed performance. Many upgrades are possible for applications requiring high imaging speed or large FOV. It is important to note that camera upgrades will likely require corresponding upgrades in the peripheral computer, data transfer links, and storage. Affordable options for both speed and FOV upgrades are also possible. The table below lists some options:

| Camera | Pixels | Speed (FPS) | Notes |
| --- | --- | --- | --- |
| Prime BSI Express | 2048 x 2048 | 95 | 95% QE |
| Kinetix | 3200 x 3200 | 500+ | 95% QE, KHz FPS, FOV |
| Iris 15 | 5056 x 2960 | 30 | Large FOV |
| pco.edge 4.2 | 2048 x 2048 | 100 | 82% QE |
| pco.panda 4.2 bi | 2048 x 2048 | 40 | 95% QE |
| pco.dicam C8 | 2048 x 2048 | 100+ | Intensified for KHz FPS |
| Blackfly S USB3 | 1600 x 1100 | 196 | Affordable CMOS, speed |
| Blackfly S GigE | 5472 x 3648 | 6 | Affordable CMOS, large FOV |

### Alternate light-sheet creation strategies

Here we have used a DSLM approach for light-sheet creation which has the advantage of shadow artifact free imaging, deep inside scattering tissues. However, this requires an additional galvanometer scanner in the setup. This approach may also be limiting in upgrading the microscope for multi-KHz frame rate imaging. An alternate strategy for light-sheet creation may involve a Powell lens combined with a converging lens. In this implementation, both the Powell lens and the corner mirror could be mounted on a common manual translation stage to introduce the required offset in the illumination beam. This enables an easy switch between centre alignment and off-axis illumination. A laser diode is another approach for light-sheet generation. A laser diode and a slit aperture can replace a standard laser and galvanometer mirror scanner combination to provide highly cost-efficient light-sheet genesis. Multicolor imaging capability in a light-weight design can also be incorporated with the recently developed matchbox formfactor lasers.

### Alternate configurations

Multiple combinations of microscope objectives are possible to support a wide range of applications. The table below lists some of these configurations, along with their key advantages. None of these configurations have been reported in the past.

| MO1 | MO2 | MO3 | System NA | Remarks |
| --- | --- | --- | --- | --- |
| 20x, 1.0 NA, Water | 20x, 1.0 NA, Water | 20x, 1.0 NA, Water | ~0.6 | > 1 mm FOV |
| 60x, 1.1 NA, Water | 60x, 1.0 NA, Water | 60x, 1.0 NA, Water | ~0.66 | Correction collar, multiphoton |
| 60x, 1.1 NA, Water | 60x, 1.1 NA, Water | 60x, 1.1 NA, Water | ~0.8 | Correction collar, multiphoton |
| 60x, 1.2 NA, Water | 60x, 1.1 NA, Water | 60x, 1.1 NA, Water | ~0.87 | Correction collar, multiphoton |
| 60x, 1.27 NA, Water | 60x, 1.1 NA, Water | 60x, 1.1 NA, Water | ~0.93 | Correction collar, multiphoton |
| No *Herschel's* configurations |  |  |  |  |
| 20x, 1.0 NA, Water | 60x, 1.0 NA, Water | 60x, 1.0 NA, Water | ~0.33 | >1 mm FOV, high collection efficiency |
| 20x, 0.5 NA, Water | 60x, 1.0 NA, Water | 60x, 1.0 NA, Water | ~0.33 | > 1 mm FOV, low cost |

*Crossbill Design* does not currently estimate NA for the no Herschel’s condition configurations. The values in the table were manually calculated by considering acceptance cone overlap and relative magnifications. The designs can also be extended with oil immersion objective lenses as MO1. Any of the existing designs with or without water immersion objectives can also be implemented on the *Crossbill* platform.

Although the demonstrated design shows OPM configuration with a SOPi scan engine, a more compact and much cheaper design is realizable by removing the SL1, G1, SL2 from the setup. This configuration is suitable for applications where samples can be easily scanned on a translation stage and a fast lateral scan is not crucial. Addition of z-axis automated control in the motorized translation stage is a convenient upgrade for facilitating 3D sample scanning.

